# Design of Parameter Intervals to Meet Steady-State Specifications in Biomolecular Circuits using Interval Analysis

**DOI:** 10.1101/2024.10.24.620126

**Authors:** Rudra Prakash, S. Janardhanan, Shaunak Sen

## Abstract

It is often desired, in the analysis or the design of biomolecular circuits, to search for parameters that satisfy certain specifications, such as on the number of the steady states or on their magnitudes. These problems are challenging because of the presence of multiple parameters, the nonlinear mapping between the parameters and the steady states, as well as the specifications themselves, such as when multiple steady states are desired. Typically, exact analytical solutions are limited and numerical approaches, if they converge, may not capture all solutions. We used Interval Analysis versions of Bisection and Constraint Propagation to obtain rigorous and guaranteed estimates of circuit parameters for the design problem. We established criteria that rule out the existence of design solutions in a given parameter space. We presented algorithms to rigorously bound all solutions and developed variants that enclosed the solutions as accurately as required. These theoretical results were illustrated on benchmark feedback and feedforward circuits. These results should aid in the analysis and design of biomolecular circuits as well as in other contexts with similar modelling frameworks. The rigorous nature of the results may be particularly useful for resource optimization and safety criticality in synthetic biology applications, such as in therapeutic applications and drug delivery.

## 1. Introduction

Mathematical models are an important component in the analysis and design workflows of biomolecular systems [1, 2, 3]. In the analysis of steady states of these models, the main emphases has been on ascertaning the qualitative dependence of the number of steady states on the parameters as well as on characterizing the quantitative robustness of the steady state on few selected parameters. Theoretical frameworks for classes of such models exist, such as those based on Chemical Reaction Network Theory [4, 5] and on Monotone Dynamical Systems [6, 7]. A much larger class of models is accessible through numerical simulations. These complement the theoretical analyses and underlie many important screens for robustness [8, 9]. Typically, these screens involve a random sampling of the parameter space to draw out conclusions on the steady-state behaviour [10, 11, 12]. However, these simulations do not generally represent rigorous numerical constructions. Recent approaches have been developed to rigorously address these questions for the dependence of the steady states on the parameters [13, 14]. The complementary question of rigorously obtaining parameters that satisfy desired steady-state specifications is an important challenge in biomolecular circuit design.

There are at least three striking aspects in designing parameters to satisfy given steady-state specifications. One is the presence of multiple parameters, which can be more than the number of steady-state constraints available. This results in an underdetermined problem that need not have a unique solution. Two is the typically nonlinear nature of the mapping from the parameters to the steady states, which may not be available in an explicit analytical form. While numerical approaches are available, these approaches may not capture all solutions, assuming they converge. Three is the specification itself, which, in biomolecular circuits, could include constraints on the number of the steady states as well as on their magnitudes. These are qualitatively different from typically assigned steady-state specifications in control engineering. Given these, methods that can rigorously identify parameters for the design of required steady states are likely to be useful.

Here we ask if, and how, given a biomolecular circuit and a design specification of desired steadystate intervals, corresponding parameter intervals may be rigorously obtained. We addressed this problem using the methods of Interval Analysis, specifically those based on Bisection and Constraint Propagation. We provided easily verifiable conditions to rule out the existence of design solutions in a given parameter box. In the case that a solution could exist, we presented algorithms to rigorous bound the solution. The solution bounds could be made as sharp as required using a subdivision of the parameter space. Steady-state specifications requiring the simultaneous existence of multiple steady states could also be solved. These results were illustrated using benchmark examples of feedforward and feedback loops.

## 2. Background

We addressed these questions using versions of Bisection and Constraint Propagation based on Interval Analysis, an approach based on set-based, rather than point-based, computations [15]. A key advantage of this approach is its ability to produce rigorous theoretical constructions. We briefly summarized the main aspects of Interval Analysis used in this work.

### 2.1 Intervals

Intervals such as 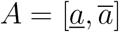 and 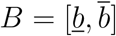, where 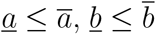, and 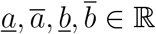 are the basic building blocks in Interval Analysis. They may be viewed as belonging to 𝕀ℝ, the set of all compact intervals of ℝ. Intervals in higher dimensions, or *boxes*, can be constructed using the cartesian product. For example, the rectangle *A × B* is an element of 𝕀ℝ^2^ = 𝕀ℝ × 𝕀ℝ. The *diameter* of an interval *A* is its width 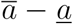 and the *radius* (rad) of *A* is one-half of the diameter. The diameter of a box may be taken as the maximum of the diameters in each dimension.

### 2.2 Interval Arithmetic

Arithmetic operations can be extended to intervals in a set-theoretic sense, for example, *A * B* = *{a * b* : *a ∈ A, b ∈ B}*, where *\* ∈ {*+, *∓, ×, ÷}*. For the *÷* operator, the above definition is for when 0 *∉ B* (When 0 *∈ B*, an extended interval division based on signed zeros may be used). An important property of this arithmetic is *inclusion isotonicity* : *A*_1_ *⊆ A*_2_ and *B*_1_ *⊆ B*_2_ *⇒A*_1_ *\* B*_1_ *⊆ A*_2_ *\* B*_2_. Interval arithmetic is the foundation for developing the notion of Interval Functions.

### 2.3 Interval Functions

Consider a rational function *f* (*x*) = *p*(*x*)*/q*(*x*), where *p*(*x*) and *q*(*x*) are polynomials. Such a function consists of a finite number of arithmetic operations and can be represented as a *directed acyclic graph* (Fig. 1 (a)). The nodes of this graph are the sub-expressions of *f* (*x*) and the edges signify how the sub-expressions combine arithmetically to build up the entire function. A *natural interval extension* of *f*, denoted *F*, is obtained by replacing all instances of the variable *x* with its interval version *X* and using interval versions of the arithmetic operations (Fig. 1 (b)). An important application of the extension *F* is in providing a means to describe the *range, R*(*f*; *X*) = *{f* (*x*) : *x ∈ X}*. The following theorem highlights two important properties of interval function extensions that help in enclosing the range.

**Figure 1:**
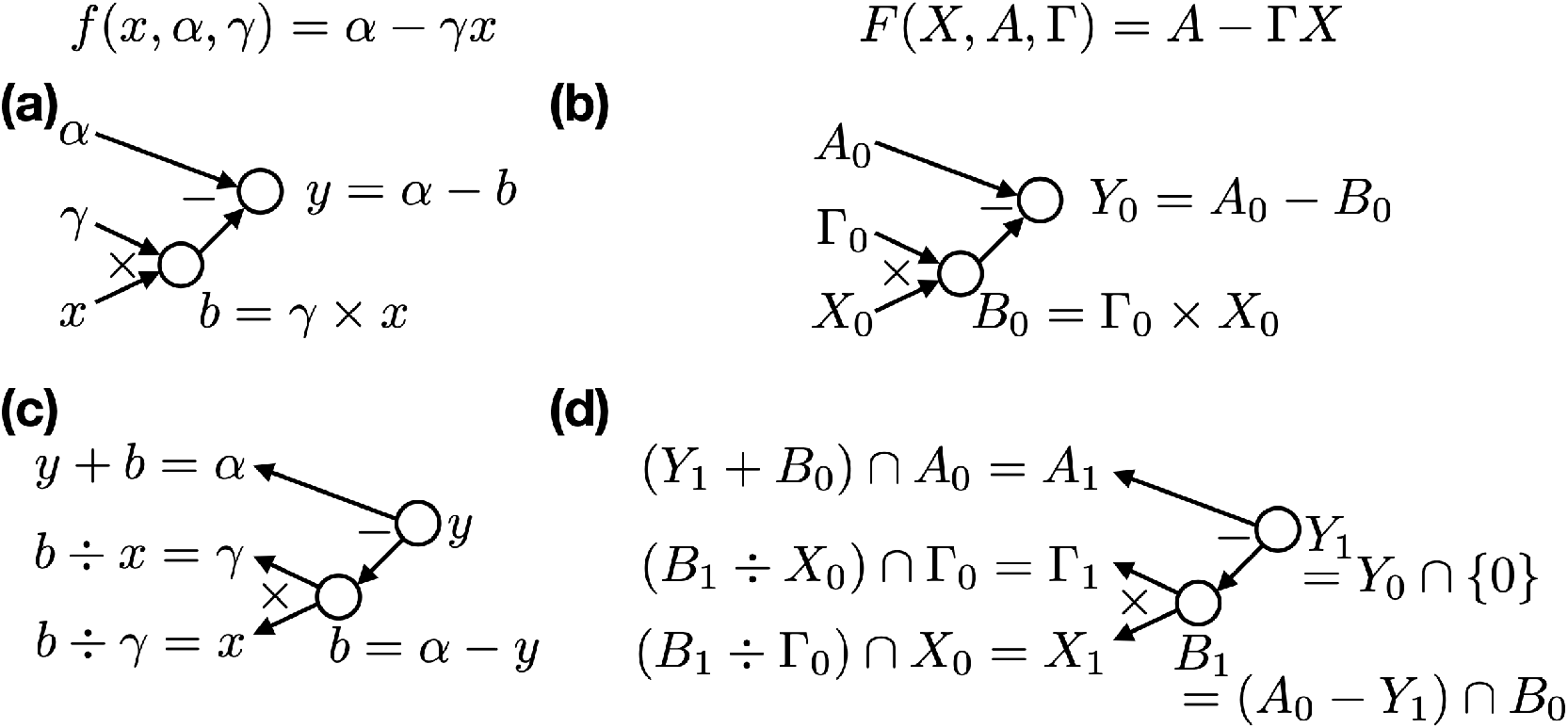
Directed Acyclic Graph (a) Nodes and edges represent the construction of *f* (*x, α, γ*). (b) Interval Extension of (a). (c) Backward direction of (a). (d) Backward direction of (b) for the constraint *f* (*x, α, γ*) = 0

#### Theorem 1.

*[15] Given a real-valued, rational function f and a natural interval extension F such that F* (*X*) *is well-defined for some X ∈* 𝕀ℝ, *we have*

1. *Y ⊆ Z ⊆ X ⇒ F* (*Y*) *⊆ F* (*Z*), *(Inclusion Isotonicity)*
2. *R*(*f*; *X*) *⊆ F* (*X*). *(Range Enclosure)*

An interval extension *F* is said to be *sharp* if *R*(*f*; *X*) = *F* (*X*). In general, the interval function extensions may not be sharp because of interval *dependency*, where a variable is encountered multiple times in the function expression. For example, for *X* = [0, 2] the natural extension of *f* (*x*) = *∓* 2*x*+*x*^2^ is *F* (*X*) = [*∓* 4, 4], but *R*(*f*; *X*) = [ *∓*1, 0] *⊂ F* (*X*). A subdivision strategy can be used to obtain sharper range enclosures. For Lipschitz continuous functions, it is guaranteed that the range can be enclosed as sharply as needed.

#### Theorem 2.

*[15]* ***(Range Enclosure)*** *Consider f* : *I → R with f Lipschitz and let F be an inclusion isotonic interval extension of f such that F* (*X*) *is well-defined for some X ⊆ I. Then there exists a positive real number k, depending on F and X, such that, if* 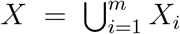 than 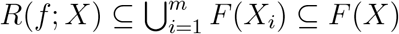 *and rad* 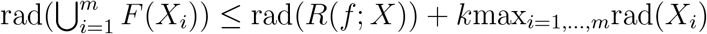.

### 2.4 Interval Bisection

The Bisection method is a classical algorithm for finding zeros of a continuous real-valued function. However, it needs a good initial guess and may not be able to locate all zeros. In its Interval version [15], [16], this method transforms into a powerful tool that circumvents these issues. Interval Bisection recursively rules out boxes that do not contain a zero by repeated application of Theorem 1: If 0 */ F* (*X*), then *X* does not contain a root of *f∈* (*x*) = 0 because *x ∈ X ⇒f* (*x*) *∈ f* (*X*) *⊆ F* (*X*). The sets that do not contain a root are discarded. The process continues till the sets that are bisected reach a pre-specified tolerance. The remaining sets *may* have a root.

### 2.5 Interval Constraint Propagation

Interval Constraint Propagation uses Interval Analysis to propagate constraints that can be expressed as functions along their Directed Acyclic Graphs in the forward and backward directions (Fig. 1, [16], [17]). The forward path evaluates the interval variables and various sub-expressions using interval arithmetic, and is essentially the natural interval extension. The backward path inverts the operations, in a set-theoretic sense, along each edge, followed by an intersection, or projection, at each node with the previous interval value [18]. Because of the projection, each cycle of forward-backward propagation *contracts* the starting interval. For a pre-specified tolerance condition, the contraction cycles stop in a finite number of steps [19]. In some instances, it has been noted that the iteration reaches a *deadlock* and does not contract further [18]. It is suggested that this be resolved by bisecting, and, in each bisected part, checking for any deadlock, and possibly redoing the constraint propagation.

## 3. Theoretical Results

### Design Problem

The steady states of a biomolecular circuit *x∈* ℝ^*n*^ are a solution to an equation of the form *f* (*x, p*) = 0, where *p ∈* ℝ^*m*^ are the parameters and *f* : ℝ^*n*^ *×* ℝ^*m*^ *→* ℝ^*n*^. Denote *X, P*, and *F* as the interval versions of *x, p*, and *f*, respectively. Given a desired steady-state interval box

*X*_*d*_ *⊆* ℝ^*n*^, find the set of parameters *S* = *{p* : *p ∈ P*_0_ *⊆* ℝ^*m*^, *∃ x ∈ X*d such that *f* (*x, p*) = 0*}* rigorously and as sharply as possible. An initial parameter interval box *P*_0_, based on biological plausibility, is assumed to be given.

The interval box *X*_*d*_ may be singleton. However, in general, both from the point of view of typical computer arithmetic and experimental measurements, a characterization of *X*_*d*_ as an interval box is well-posed.

The interval *X*_*d*_ may also be a set of disjoint interval boxes. These specifications often arise in biomolecular circuits, both in developmental biology as well as in the design of switches. The following theorem provides a way to solve such specifications.

#### Theorem 3.

***(Multiple Steady States)*** *Suppose X*_*d*_ = *X*_*d*1_ *∪ X*_*d*2_ *where X*_*d*1_ *∩ X*_*d*2_ = *∅. Then the solution to the design problem exists in S*_1_ *∩ S*_2_, *where S*_1_ *and S*_2_ *are solution enclosures of X*_*d*1_ *and X*_*d*2_, *respectively*.

*Proof*. Both solutions lie in *S*_1_ *∩ S*_2_. ▄

Typical models based on mass action give rise to functions *f* that are smooth. Often, piecewise linear approximations are used, where *f* is not differentiable over the whole domain. We assume that *f* is Lipschitz continuous.

The bounded nature of the states and parameters and the continuity properties of the steady-state equation naturally lend themselves to solution approaches based on Interval Analysis.

### 3.1 Interval Bisection

We first state a non-existence criterion based on the Interval version of Bisection.

#### Theorem 4.

***(Non-existence Criterion 1)*** *The design problem does not have a solution in P*_0_ *if* 0 *∈/ F* (*X*_*d*_, *P*_0_).

*Proof*. The proof is by contradiction. Suppose 0 *∈ F* (*X*_*d*_, *P*_0_) and the design problem has a solution in *P*_0_. The existence of a solution *⇒ ∀x ∈ X*_*d*_ *∃ p ∈ P*_0_ such that *f* (*x, p*) = 0. From the Inclusion Isotonicity property in Theorem 1, (*x, p*) *∈ X*_*d*_ *×P*_0_ *⇒ f* (*x, p*) *∈ f* (*X*_*d*_, *P*_0_). From the Range Enclosure property in Theorem 1, *f* (*X*_*d*_, *P*_0_) *⊆ F* (*X*_*d*_, *P*_0_). Therefore, 0 = *f* (*x, p*) *∈ f* (*X*_*d*_, *P*_0_) *⊆ F* (*X*_*d*_, *P*_0_).

This is a contradiction. ▄

**Corollary 1**. *If the interval extension F of f is sharp, then the condition* 0 *∈ F* (*X*_*d*_, *P*_0_) *is necessary and sufficient for the solution to exist in P*_0_.

*Proof*. If *F* is a sharp interval extension of *f*, then the Range Enclosure property ensures an equality *f* (*X*_*d*_, *P*_0_) = *F* (*X*_*d*_, *P*_0_). The result follows. ▄

This directly gives rise to algorithms for zeroing on the desired parameter intervals.

#### Theorem 5.

***(Recursive Bisection)*** *The solution, if it exists in P*_0_, *can be bounded by Algorithm 1*.

#### Algorithm 1: Recursive Bisection

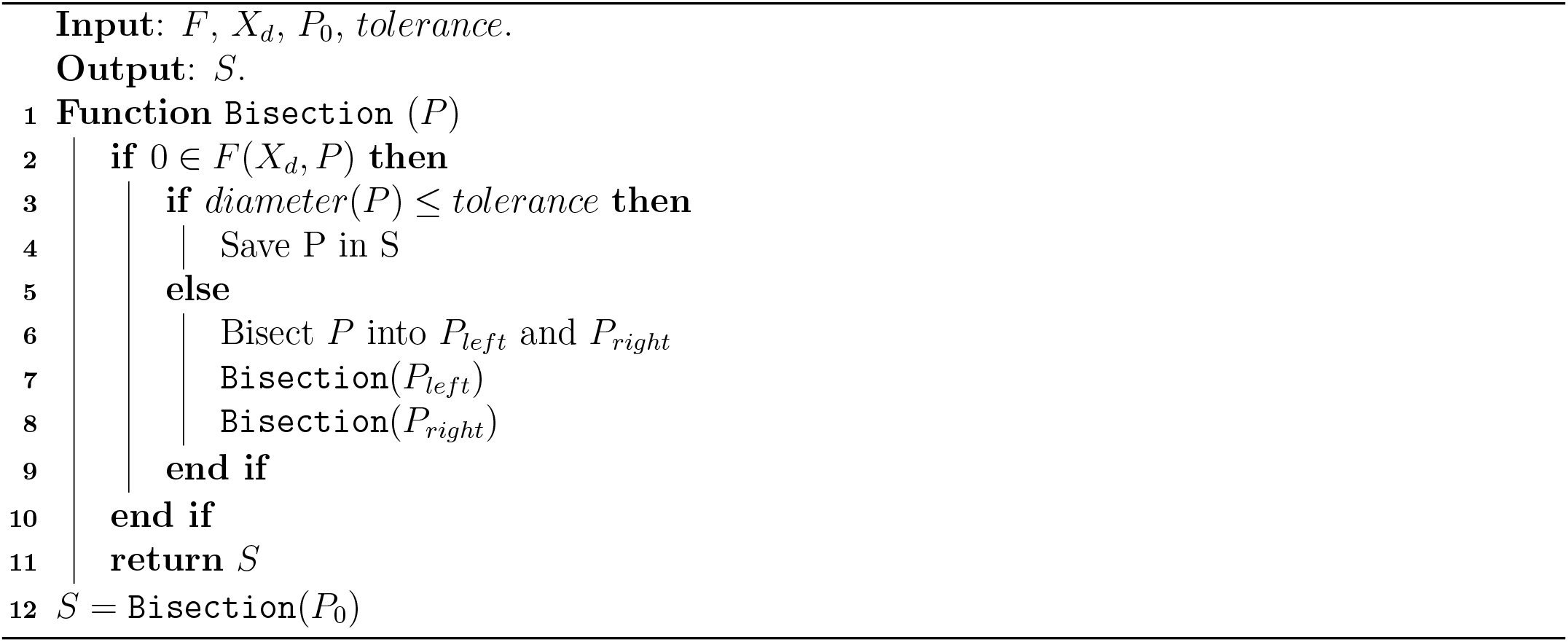

*Proof*. The algorithm recursively bisects *P*_0_ and discards intervals that do not contain 0. The bisections are continued until the diameter of the bisected set is smaller than a pre-specified tolerance. Because the regions that do not contain a solution are discarded, the remaining region must have a solution, if the solution exists in *P*_0_. As the diameter necessarily decreases with each bisection, this algorithm converges in a finite number of steps. ▄

We note that this bisection algorithm is not adaptive in the sense that the size of the final sets is solely determined by the tolerance condition. This arises because the design problem is in the form of an equality constraint and not an inequality constraint that can be treated as an inclusion.

A standard drawback of recursion-based algorithms is the multiple number of times each region may need to be evaluated. This can be circumvented by subdividing the parameter space into subintervals and then filtering the subintervals using the Non-Existence Criterion of Theorem 4.

#### Theorem 6.

***(Subdivision*** + ***Filter)*** *The solution, if it exists in P*_0_, *can be bounded as sharply as needed by combining parameter subdivision with a filter (Algorithm 2)*.

#### Algorithm 2: Subdivision + Filter

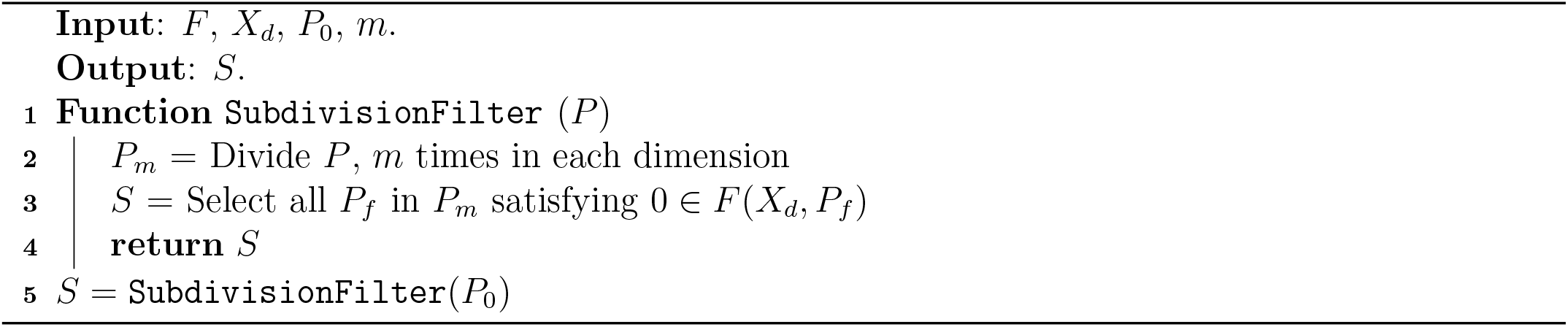

*Proof*. Given parameter interval *P*_0_ is subdivided into *m* intervals in each direction based on the sharpness required. The subintervals *P*_*f*_ whose map *F* (*X*_*d*_, *P*_*f*_) does not contain 0 are discarded. The remaining subintervals must have a solution, if it exists in *P*_0_. Decreasing the width of the subintervals increases the sharpness of the solution enclosure. ▄

The advantage of the recursive bisection algorithm, of ruling out large regions in the parameter space without the need for checking the Non-Existence Criterion for each subinterval, can be incorporated in the subdivision algorithm.

#### Theorem 7.

*The solution, if it exists in P*_0_, *can be bounded as sharply as required by an initial recursive bisection followed by parameter subdivision with a filter*.

*Proof*. A recursive bisection, without any tolerance condition, initiates the search and discards regions that do not have a solution. When both halves of the set have a solution, the set is subdivided and its subintervals are filtered. The width of the subintervals determines the sharpness. ▄

We noted that the design problem posed is a many-to-one problem in that multiple parameters could give the same steady state. The formulation of the above algorithms is not limited by the number of parameters. Therefore, they apply naturally for any number of parameters *m ≥* 1.

### 3.2 Interval Constraint Propagation

The *n* equations in the design problem *f* (*x, p*) = 0 for the desired steady-state interval may be viewed as constraints. Each function *f* can be represented as a Directed Acyclic Graph with the parameters *p* and the steady-state variables *x* as the inputs and the function evaluation *f* (*x, p*) as the output. For typical biomolecular circuits, each intermediate node *c* in the graph represents the endpoint of an operation of the form *a * b* = *c*, where *a* and *b* are the nodes or inputs, connected to *c* and *\*∈ {*+,*∓, ×, ÷*, ^*}*. The operations + and *∓* arise when the rates of two reactions combine. The operation *×*arises in reaction rates when two states or a state and a parameter are multiplied. The operation *÷* arises when a ratio of reaction rates is considered, for example, in obtaining a dissociation constant from the forward and backward reaction rates. The operation ^ arises in terms containing the Hill coefficient.

Interval versions of the parameters are propagated along the graph in the forward direction. For this, interval versions of arithmetic operations are performed at each node. The propagation of the initial parameter interval *P*_0_, for the desired steady-state interval *X*_*d*_, yields *F* (*X*_*d*_, *P*_0_). The steadystate constraint *f* (*x, p*) = 0 is applied as *F* (*X*_*d*_, *P*_0_)*∩{*0*}* and the resulting set (*{*0*}* or *∅*) is propagated in the backward direction. If an operation *A*_0_ *\* B*_0_ = *C*_0_ is performed at a node *c* in the forward direction, the corresponding inverse operations are to be performed and propagated to the connecting nodes *a* and *b* in the backward direction as *A*_1_ = (*C*_1_ *\**^*∓*1^ *B*_0_) *∩ A*_0_ and *B*_1_ = (*C*_1_ *\**^*∓*1^ *A*_0_) *∩ B*_0_, where *\**^*∓*1^ is the appropriate inverse operation of *\** and the subscripts 0 and 1 represent the initial and final versions of the node values (*A, B, C*), respectively. In the case that a parameter appears at multiple nodes in the Directed Acyclic Graph, the backward paths are evaluated in parallel and an intersection is taken. The backward propagation results in a parameter interval *P*_1_, which is a subset of *P*_0_, because the intervals obtained at each node in the backward direction are a subset of the corresponding intervals in the forward direction, for example, *A*_1_ *⊆ A*_0_ and *B*_1_ *⊆ B*_0_. One cycle of this forward-backward propagation *C* : *P*_0_ *→ P*_1_ is referred to as the *contractor*.

The following Non-Existence Criterion can be stated.

#### Theorem 8.

***(Non-existence Criterion*** 2) *The design problem does not have a solution in P*_0_ *if C*(*P*_0_) = *∅*.

*Proof*. The proof is by contradiction. Suppose *C*(*P*_0_) = *∅* and the design problem has a solution in *P*_0_. The existence of a solution *⇒ ∀ x ∈ X*_*d*_ *∃ p ∈ P*_0_ such that *f* (*x, p*) = 0. (*x, p*) satisfies each node of the forward and backward path of the Directed Acyclic Graph. Therefore, *C*(*P*_0_) ≠ *∅*. This is a contradiction. ▄

This also shows that *p∈ S ⊆ C*(*P*_0_) if *S∩ P*_0_ ≠ *∅*.

The Non-Existence criterion in Theorem 8 is stronger than the Non-Existence Criterion in Theorem 4.

#### Theorem 9.

*C*(*P*_0_) *≠ ∅ ⇒* 0 *∈ F* (*X*_*d*_, *P*_0_).

*Proof*. Suppose 0 *∉ F* (*X*_*d*_, *P*_0_). *⇒* the backward path propagates an empty set. *⇒ C*(*P*_0_) = *∅*.

Therefore, *C*(*P*_0_) *≠ ∅ ⇒* 0 *∈ F* (*X*_*d*_, *P*_0_). ▄

The basic algorithm is to iterate the contractor.

#### Theorem 10.

***(Contractor)*** *The design solution, if it exists in P*_0_, *can be bounded by Algorithm 3*.

#### Algorithm 3: Contractor

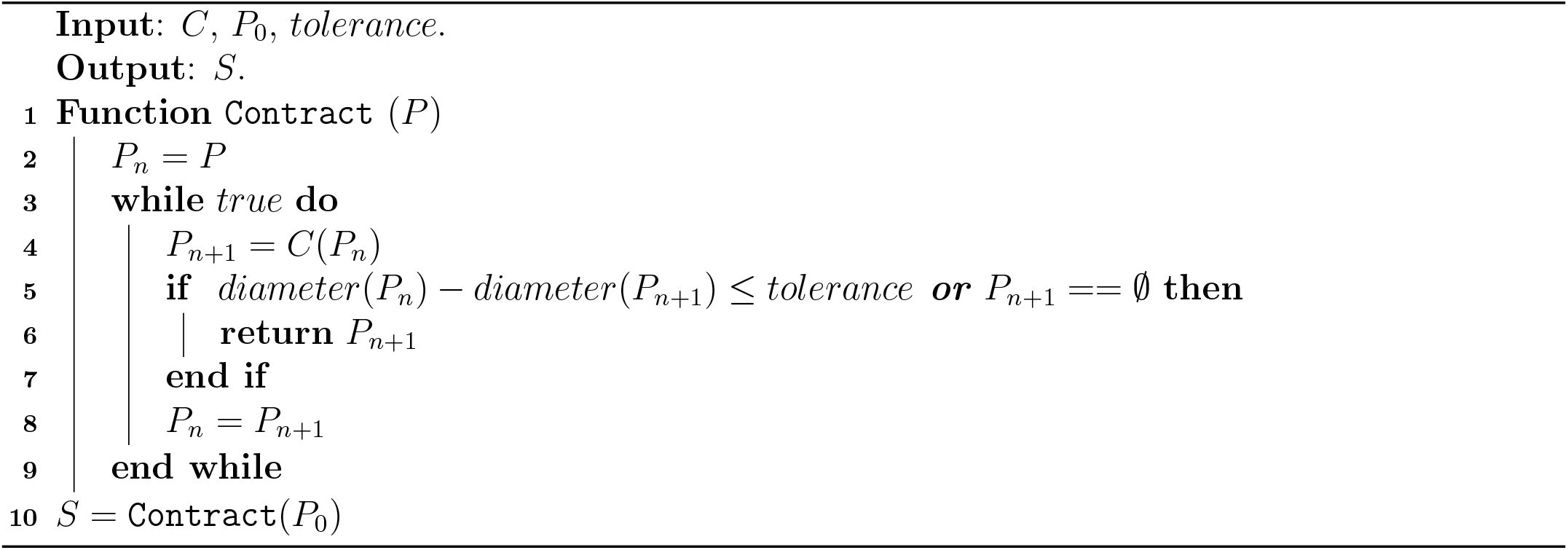

*Proof*. Each iteration of the contractor contracts the possible solution set and remains a bound for the overall solution set. The algorithm converges in a finite number of iterations because *P*_0_ is bounded. ▄

It is possible that the solution bound obtained is not sharp because a cartesian product of the individually contracted parameter intervals is taken. A variation to the Algorithm 3 is to subdivide the parameter intervals and to apply the contractor on each subdivided interval.

#### Theorem 11.

***(Subdivision*** + ***Contractor)*** *The solution, if it exists in P*_0_, *can be enclosed as sharply as needed by contracting subdivided parameter intervals (Algorithm 4)*.

#### Algorithm 4: Subdivision + Contractor

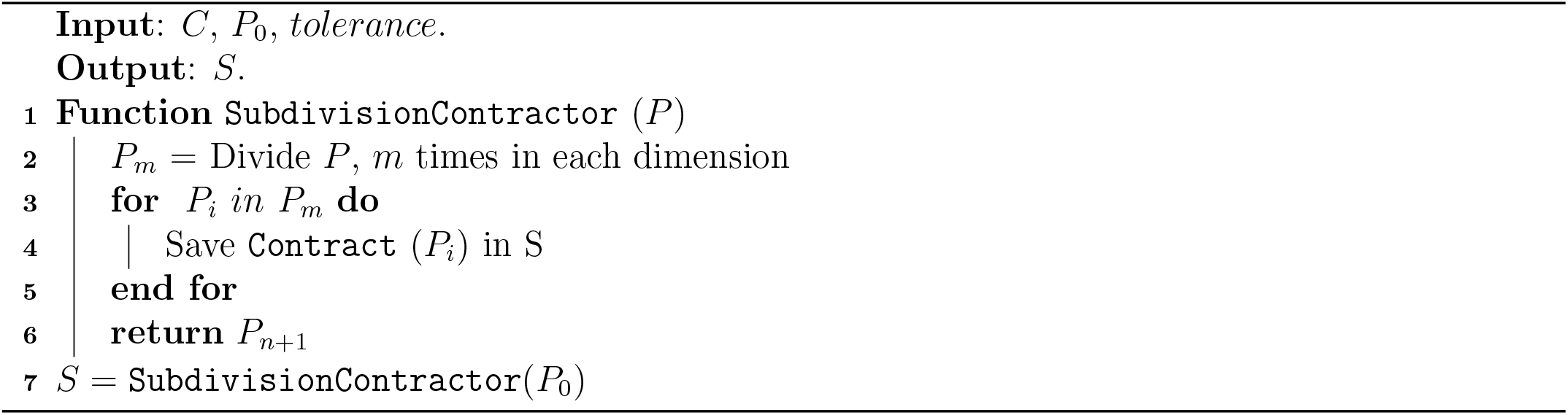

*Proof*. Subdivision reduces the overestimation as the cartesian product of subdivided individual parameter intervals is taken. The width of the subintervals can be tuned to modulate the sharpness of the bound. Essentially, the subdivision sharpens the contractor function. ▄

The advantages of the Bisection method can be incorporated into the Contractor-based approach. The Non-Existence Criterion in Theorem 4 can be used as an additional Filter in Algorithm 4 so that only those subdivided intervals that may have a solution are contracted. This Algorithm 4F may save computational costs further. The proof can be obtained by combining the proofs of Theorem 4 and Theorem 7.

A judicious combination of the above Bisection, Contractor, and Subdivision algorithms that incorporates their relative strengths can provide sharp solution enclosures at a reduced computational cost. Bisection can rule out large parts of the parameter space where there is no solution. Subdivision can reduce interval dependencies leading to sharper solution enclosures and avoid the computational costs associated with the recursion inherent in Bisection. Contractor can discard parts of the parameter space without a solution less conservatively than Bisection. Combinations of these algorithms can also resolve situations of *deadlock*, where the solution enclosure gets stuck. One approach to resolve deadlock is to subdivide in a sufficiently fine manner. Another approach is to combine Bisection and Contractor in an adaptive way (Algorithm 5). The proof follows from the the proofs of the constituent parts noted above.

#### Algorithm 5: Contractor + Bisection

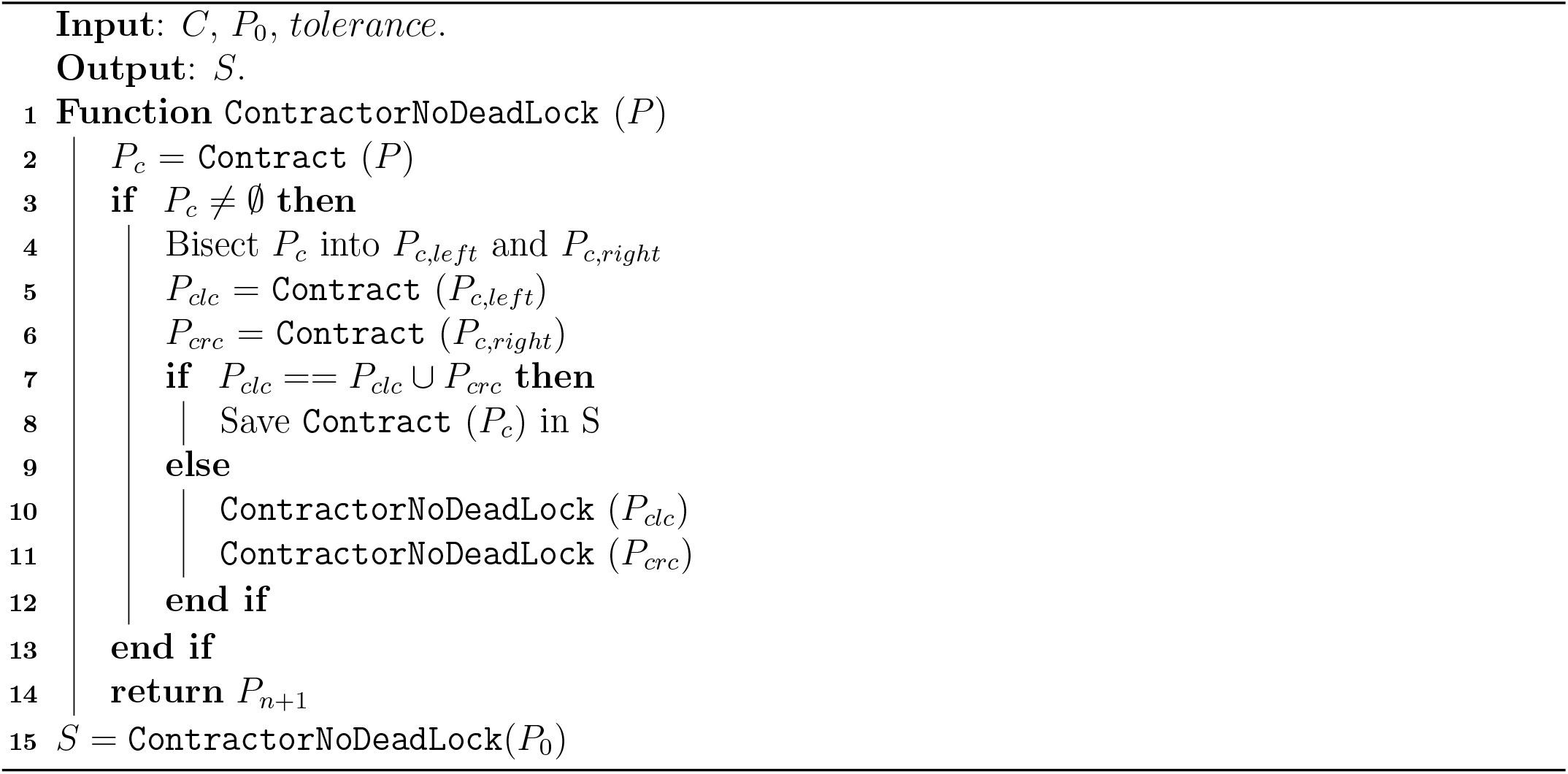

## 4. Examples

We illustrated various facets of the above methods using canonical examples of biomolecular circuits ([1], [2]). The computations were implemented in Julia 1.8.3 with the package IntervalArithmetic v0.20.9.

### 4.2 Example 1 (Open Loop)

This is a basic model for the production and degradation of a protein (Fig. 2(a), inset), chosen so that the results from the proposed algorithms can be checked against the exact solutions. The steady-state equation is *f* (*x, α, γ*) = *α ∓ γx* = 0. For *X*_*d*_ = [60, 70], *A*_0_ = [20, 30], Γ_0_ = [1, 2], *F* (*X*_*d*_, *A*_0_, Γ_0_) = [*∓*120, *∓*30]. As 0 *∉ F* (*X*_*d*_, *A*_0_, Γ_0_), it is guaranteed that the solution does not exist. For *X*_*d*_ = [20, 30], *F* (*X*_*d*_, *A*_0_, Γ_0_) = [ − 40, 10] 0 and a solution may exist. This conforms to the expectations from the exact solution.

**Figure 2:**
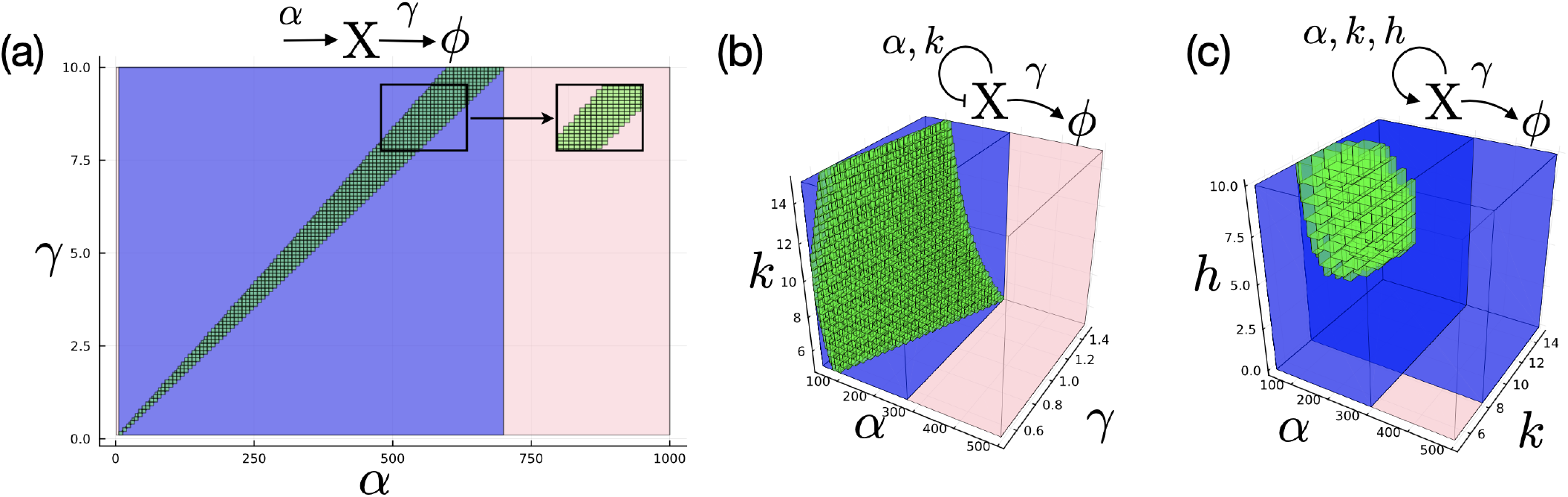
Examples: Pink, blue, and green boxes represent the initial parameter box (*P*_0_), the contractor after one iteration (*P*_1_, where applicable), and the final solution enclosure (*S*), respectively. Circuit schematics are shown above the plots. (a) Example 1 : *P*_0_ = *A*_0_ *×* Γ_0_ = [1, 1000] *×* [0.1, 10]. The inset shows the indicated portion of the solution enclosure using Algorithm 2. (b) Example 2: *P*_0_ = *A*_0_ *×* Γ_0_ *× K*_0_ = [50, 500] *×* [0.5, 1.5] *×* [5, 15]. (c) Example 3: *P*_0_ = *A*_0_ *× K*_0_ *× H*_0_ = [50, 500] *×* [5, 15] *×* [0, 10]. Γ = [0.5, 1.5] has been fixed for ease of visualization. Green boxes represent *S*_1_ *∩ S*_2_, where *S*_1_ and *S*_2_ are the solution enclosures for *X*_*d*1_ and *X*_*d*2_, respectively. Blue boxes are the respective enclosures to the solutions after one contractor iteration.

The enclosure of the solution using Algorithm 4F (Algorithm 4 + Filter based on Non-Existence Criterion 1) with *X*_*d*_ = [60, 70] and *m* = 100 subdivisions in each parameter dimension is shown in Fig. 2(a). The forward path in each contractor cycle (Fig. 1) is *B*_0_ = Γ_0_ *× X*_*d*_, *Y*_0_ = *A*_0_ *∓ B*_0_. Then, the steady-state constraint *Y*_1_ = *Y*_0_ *∩ {*0*}* = *{*0*}* is applied, followed by the backward path *A*_1_ = (*B*_0_ + *Y*_1_) *∩ A*_0_, *B*_1_ = (*A*_0_ *∓ Y*_1_) *∩ B*_0_, Γ_1_ = (*B*_1_*/X*_*d*_) *∩* Γ_0_. For the same *X*_*d*_ and *m*, a portion of the solution enclosure using Algorithm 2 is shown in the inset of Fig. 2(a). The boundaries of the solution enclosure in Fig. 2(a) were smoother than those in the inset. This was because all boxes in the inset were of the same size, whereas the contractor can make the boxes smaller.

### 4.2 Example 2 (Negative feedback)

In this model, the protein negatively regulates its own expression, forming a negative feedback (Fig. 2(b), inset). The steady-state equation is *f* (*x, α, γ, k*) = *α/*(1 + *x/k*) *∓γx* = 0. As a network motif, this circuit is a building block of biomolecular circuits [2].

For *X*_*d*_ = [26, 28] and 25 subdivisions in each parameter dimension, an enclosure of the solution using Algorithm 4F is shown in Fig. 2(b). The forward path in each contractor cycle is *B*_0_ = Γ_0_ *× X*_*d*_, *C*_0_ = *X*_*d*_*/K*_0_, *D*_0_ = 1 + *C*_0_, *E*_0_ = *A*_0_*/D*_0_, *Y*_0_ = *E*_0_ *∓ B*_0_. Following the steady-state constraint *Y*_1_ = *Y*_0_ *∩* 0 = 0, the backward path *E*_1_ = (*B*_0_ + *Y*_1_) *∩ E*_0_, *B*_1_ = (*E*_0_ *∓ Y*_1_) *∩ B*_0_, *A*_1_ = (*E*_1_ *× D*_0_) *∩ A*_0_, *D*_1_ = (*A*_0_*/E*_1_) *∩ D*_0_, *C*_1_ = (*D*_1_ *∓* 1) *∩ C*_0_, *K*_1_ = (*X*_*d*_*/C*_1_) *∩ K*_0_, Γ_1_ = (*B*_1_*/X*_*d*_) *∩* Γ_0_ is computed.

An exact analytical solution of the steady-state equation in terms of a quadratic equation would also give a solution surface in the parameter space for each point in the desired steady-state interval *X*_*d*_. However, it would have to be calculated numerically, possibly for a random sampling in *X*_*d*_. For variations of this model, when an exact solution may not be available or provide insight, such as when a Hill co-efficient *h* is present in the production term *α/*(1 + (*x/k*)^*h*^), only a numerical approach may be possible. In contrast to this, the solution enclosure computed above is rigorous and can be made as sharp as required.

### 4.3 Example 3 (Positive feedback)

In this model, the protein can activate its own expression, resulting in a positive feedback. The steady-state equation is *f* (*x, α, γ, k*) = *α/*(1 + (*k/x*)^*h*) *∓ γx* = 0. This is a widely used model for contexts with multiple steady states such as switches and development. Therefore, we chose a steady-state specification that had disjoint intervals *X*_*d*1_ = [7, 8] and *X*_*d*2_ = [100, 101].

An enclosure of the solution using Algorithm 4F with *m* = 10 subdivisions is shown in Fig. 2(c). The forward path in each contractor cycle is *B*_0_ = *K*_0_ *÷ X*_*d*_, *C*_0_ = *B*_0_^*H*_0_, *D*_0_ = 1 + *C*_0_, *E*_0_ = *A*_0_ *÷ D*_0_, *F*_0_ = Γ *× X*_*d*_, *Y*_0_ = *E*_0_ *∓ F*_0_. Then, the steady-state constraint *Y*_1_ = *Y*_0_ *∩* 0 = 0 was applied. Finally, the backward path is applied *F*_1_ = (*E*_0_ *∓ Y*_1_) *∩ F*_0_, *E*_1_ = (*Y*_1_ + *F*_0_) *∩ E*_0_, *D*_1_ = (*A*_0_ *÷ E*_1_) *∩ D*_0_, *A*_1_ = (*E*_1_ *× D*_0_) *∩ A*_0_, *C*_1_ = (*D*_1_ *∓* 1) *∩ C*_0_, *B*_1_ = (*C*_1_^(1*/H*_0_)) *∩ B*_0_, *H*_1_ = (*logC*_1_ *÷ logB*_0_) *∩ H*_0_, *K*_1_ = (*B*_1_ *× X*_*d*_) *∩ K*_0_.

It is frequently required, in analysis and design contexts, to find the region in parameter space that allows for the existence of multiple steady states. The proposed approach provides a conceptually simple and practically rigorous way to demarcate this region. This is in addition to the advantages over numerical methods in computing the exact solution, assuming an analytical expression can be obtained.

### 4.4 Example 4 (Incoherent Feedforward Loop)

The incoherent feedforward loop is an important example of a biomolecular circuit exhibiting adaptation. It also belongs to the class of feedforward loops identified as network motifs [2]. The steady-state equation of a simple incoherent feedforward loop is *f* (*x, y, α, γ, k*) = [*α∓ γx, α/*(1 + *x/k*) *∓ γy*]^*T*^ = [0, 0]^*T*^ .

This example has two variables. It also has parameter variables like *α* and *γ* that appear in two places. Therefore, the constraints in the backward path are evaluated in parallel and an intersection is taken.

The following steps may be used to compute a solution enclosure using Algorithm 4F. The forward path in each contractor cycle is *B*_0_ = *X*_*d*_ *÷ K*_0_, *C*_0_ = 1 + *B*_0_, *D*_0_ = Γ_01_ *× X*_*d*_, *E*_0_ = Γ_02_ *× Y*_*d*_, *F*_0_ = *A*_02_ *÷ C*_0_, *Z*_1_ = *A*_01_ *∓ D*_0_, *Z*_2_ = *F*_0_ *∓ E*_0_ where Γ_01_ = Γ_0_ = Γ_02_ and *A*_01_ = *A*_0_ = *A*_02_. After the steady-state constraints *Z*_1_ = *Z*_1_ *∩* 0 = 0, *Z*_2_ = *Z*_2_ *∩* 0 = 0, the backward path was computed using *F*_1_ = (*Z*_2_ + *E*_0_) *∩ F*_0_, *E*_1_ = (*F*_0_ *∓ Z*_2_) *∩ E*_0_, *D*_1_ = (*A*_01_ *∓ Z*_1_) *∩ D*_0_, *A*_11_ = (*D*_0_ + *Z*_1_) *∩ A*_01_, *C*_1_ = (*A*_02_ *÷ F*_1_) *∩ C*_0_, *A*_12_ = (*C*_0_ *× F*_1_) *∩ A*_02_, Γ_12_ = (*E*_1_ *÷ Y*_*d*_) *∩* Γ_02_, Γ_11_ = (*D*_1_ *÷ X*_*d*_) *∩* Γ_01_, *B*_1_ = (*C*_1_ *∓* 1) *∩ B*_0_, *K*_1_ = (*X*_*d*_ *÷ B*_1_) *∩ K*_0_, *A*_1_ = *A*_11_ *∩ A*_12_, Γ_1_ = Γ_11_ *∩* Γ_12_. This example demonstrates that the proposed methods are not necessarily restricted to scalar examples and may be formulated for circuits with multiple variables as well.

## 5. Conclusions

An important design problem in a biomolecular circuit is the rigorous determination of parameters that satisfy given steady-state specifications. Here, we addressed this using Interval versions of Bisection and Constraint Propagation. We established criteria for the non-existence of a solution to the design problem. When the solution existed, we presented algorithms to rigorously bound the required parameter intervals. These parameter intervals could be made as sharp as desired. We showed that specifications with multiple steady states could also be solved. We illustrated our results using case studies of feedforward and feedback loops. These results contribute to developing a rigorous framework for biomolecular circuit design, providing an important link between theoretical frameworks and numerical simulations.

Future work includes a quantitation of the algorithmic cost, in terms of the computational memory and time, as a function of the sharpness required and the system dimension as well as an expansion of the specifications to include other functionally important aspects such as the stability of the steady state and the local transient response.

Because biomolecular circuit models are similar to those in ecology and opinion dynamics, the developed framework should be helpful in those contexts as well. Preliminary work for these contexts is presented below.

### 5.1 Ecological Dynamics

One class of models used for ecological predator-prey models are the Lotka-Volterra equations [20]. Such models are the archetype of competition, and are also used in systems biology contexts of interacting populations, such as in [21]. Consider the following version of the Lotka-Volterra equations,

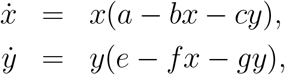

where the states *{x, y}* model the species populations and the positive parameters *{a, b, c, e, f, g}* represent the interaction terms. Typically, a geometric phase-plane analysis is performed to qualitatively determine the steady states. It is clear from inspection that (0, 0) is one steady state and the other steady-state value follows from the solution, if it exists, of a linear system of equations *a ∓bx ∓cy* = 0, *e ∓fx ∓gy* = 0. These exact solutions may be used to determine regions in parameter space that give the desired steady-state intervals. This is similar to Example 1 above.

Methods based on interval analysis may also be used to determine the regions of parameter space that give desired steady-state values. The conditions for a steady state, 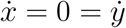, give the constraint equations. For a given steady-state specification, say *X*_*d*_ = [9, 11] *×* [4.5, 5.5], it is desired to find parameter box(es). One way to solve this problem is to use Algorithm 4. For an initial parameter interval box *P*_0_ = (*a, b, c, e, f, g*) = [0.1, 50]^6^, the contracted parameter interval box can be obtained as *P*_*c*_ = [1.35, 50] *×* [0.1, 5.51] *×* [0.1, 10.9] *×* [1.35, 50] × [0.1, 5.51] *×* [0.1, 10.9]. Subdivision of parameters could be done to make the solution enclosures sharper. Variants of this model to accommodate experimental results are relatively more complicated, for example,

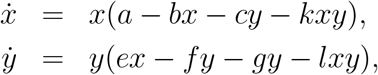

where the *xy* term represents an inhibitory effect. A quantitative determination of the steady-state values in this case is not as simple as above. For the same steady-state specification as above, an initial parameter interval box *P*_0_ = (*a, b, c, k, e, f, g, l*) = [0.1, 50]^8^, the contracted parameter interval box can be obtained as *P*_*c*_ = [5.4, 50] *×* [0.1, 5.06] *×* [0.1, 10] *×* [0.1, 1.2] *×* [5.4, 50] *×* [0.1, 5.06] *×* [0.1, 10] *×* [0.1, 1.2]. As above, subdivisions could be used to get a sharper enclosure.

### 5.2 Opinion Dynamics

Models used in opinion dynamics, for example in [22], have a similar form,

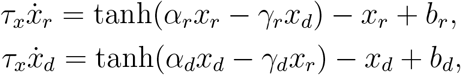

where the states *{x*_*r*_, *x*_*d*_*}* represent populations and the parameters *{α*_*r*_, *α*_*d*_, *γ*_*r*_, *γ*_*d*_, *b*_*r*_, *b*_*d*_*}* represent the interaction strengths. In an analysis in the study, *α*_*r*_ = *α*_*d*_ = *α* and *γ*_*r*_ = *γ*_*d*_ = *γ* were used. The tanh function is not a rational function but mimics the behaviour of rational functions seen in biological and ecological models in terms of being continuous, differentiable, and bounded. The steady-state constraints can be formulated using interval analysis to find the desired parameters, if they exist, given steady-state specifications. In particular for desired steadystate interval [0.475, 0.525] *×* [ *∓*0.525, *∓*0.475], a fixed *τ*_*x*_ = 5, and an initial parameter box of [1.01, 10] *×* [0.01, 1] *×* [0.001, *∓*0.1] *×* [*∓* 0.1, *∓*0.001], the contracted parameter box is [1.01, 1.21] × [0.01, 0.215] *×* [0.001, 0.075] *×* [ *∓*0.075, 0.001].

These examples provide preliminary steps towards the analysis of models from ecology and opinion dynamics. A systematic exploration of these domains would necessitate a consideration of a larger set of models.

## Acknowledgment

The authors thank Prof. I. N. Kar for his valuable feedback.

